# Life-history dynamics: damping time, demographic dispersion and generation time

**DOI:** 10.1101/2020.12.09.417261

**Authors:** Sha Jiang, Harman Jaggi, Wenyun Zuo, Madan K. Oli, Jean-Michel Gaillard, Shripad Tuljapurkar

**Author notes:** equal contribution (Sha Jiang, Harman Jaggi).

## Abstract

Transient dynamics are crucial for understanding ecological and life-history dynamics. In this study, we analyze damping time, the time taken by a population to converge to a stable (st)age structure following a perturbation, for over 600 species of animals and plants. We expected damping time to be associated with both generation time *T_c_* and demographic dispersion *σ* based on previous theoretical work. Surprisingly, we find that damping time (calculated from the population projection matrix) is approximately proportional to *T_c_* across taxa on the log-log scale, regardless of *σ*. The result suggests that species at the slow end of fast-slow continuum (characterized with long generation time, late maturity, low fecundity) are more vulnerable to external disturbances as they take more time to recover compared to species with fast life-histories. The finding on damping time led us to next examine the relationship between generation time and demographic dispersion. Our result reveals that the two life-history variables are positively correlated on a log-log scale across taxa, implying long generation time promotes demographic dispersion in reproductive events. Finally, we discuss our results in the context of metabolic theory and contribute to existing allometric scaling relationships.

## Main

In a constant environment, (st)age-structured populations tend towards a stable demographic structure (Lotka 1939; Leslie 1945, 1948; Lefkovitch 1965; Caswell 2001). However, disturbances such as environmental fluctuations, disease, and biological invasions can alter the size or structure of a population, so understanding the response to fluctuations and resulting transient dynamics are important to many questions in ecology and evolution (Lande et al. 2003; Gamelon et al. 2014). In an era when anthropogenic activities are altering global biodiversity, reshaping populations, and even driving species to extinction (Faurby and Svenning 2015) it is even more important to ask: What is the effect of external disturbances on populations? Are some species or taxa more vulnerable to perturbations than others? To help answer such questions, we analyzed the biological determinants of the damped response of populations as they recover from an external disturbance.

A population structure that experiences a disturbance returns towards stability in a sequence of damped cycles. The damping rate *d >* 0 measures how a population responds to disturbance, by quantifying the speed at which population converges to a stable (st)age distribution. The damping time *τ=* (1/*d*) is the time scale over which the effect of a disturbance dies away. A short damping time (equivalently, a large damping rate) means that population’s structure recovers rapidly from a perturbation, and implies a short demographic memory (Caswell 2001; Keyfitz and Caswell 2005). It is well known that the period of the damped cycles is close to the cohort generation time *T_c_*, the average age of survival-weighted reproduction (Keyfitz 1968). Biologically, the generation time *T_c_* measures the “pace” of life (Gaillard et al. 1989).

Earlier work found that damping time increases with generation time *T_c_* (Keyfitz 1968; Hughes and Tanner 2000) but decreases with the dispersion of reproductive events across the lifetime, measured by the demographic dispersion (Coale 1972; Taylor 1979; Keyfitz 1965; Trussell et al. 1977; Wachter 1991). Demographic dispersion *σ* is calculated as the standard deviation of survival-weighted reproduction and can be interpreted as a measure of the variation of age at reproduction around *T_c_*. Mathematically, damping time has been shown to be approximately proportional to *T_c_*^3^/*σ*^2^ (Keyfitz 1965; Keyfitz 1968). Consistent with these analyses, Coale (1972) found that the damping time in human populations decreases when reproduction is spread symmetrically over an increasing range of ages (i.e., with increasing *σ*). Taylor (1979) examined insect populations and found that changes in the age of first reproduction and demographic dispersion influence the damping time. Hughes and Tanner (2000) found that slow-growing corals with large *T_c_* have a longer damping time than fast-growing corals. However, these findings focus on groups of species (humans, insects and corals), each with a limited range of generation times. Comparative studies across taxonomic groups are needed to identify how damping time varies over a wide range of generation times.

For a large collection of age-structured data on 111 diverse mammals, Gamelon et al. (2014) examined metrics of transient dynamics and found that short-term demographic responses to disturbance are shaped by both generation time and growth rate. This study found that species characterized by long generation time and low fecundity tend to decrease in population size following a disturbance. They also conclude that these slow-living species might be more vulnerable as they are not expected to counterbalance the negative effects of disturbances by increasing population growth.

Here we use a more extensive dataset of plants and animals to analyse transient dynamics as measured by the damping time (*τ*). According to the approximation (*τ α T_c_*^3^/*σ*^2^), damping time is shaped by the two life history measures of generation time and demographic dispersion. We expected to find that damping time is proportional to the ratio *T_c_*^3^/*σ*^2^, in accordance with theory (Keyfitz and Caswell 2005). Surprisingly, we find the simple relationship that damping time is proportional to *T_c_* across taxa on the loglog scale, regardless of *σ*. Although the relationship is noisy, our result implies that time to convergence increases with generation time. In the context of the fast-slow continuum (Stearns 1983; Oli 2004; Gamelon et al. 2016), species at the slow end of the spectrum (characterized with long generation time, late maturity, low fecundity) are more vulnerable to external disturbances as they take more time to recover compared to species with fast life-histories (characterized by short generation time, high fertility, etc). This is consistent with recent work that studied demographic resilience and recovery time from disturbances (Capdevila et al. 2020; Lebreton et al. 2012).

Our result on damping time led us to examine the relationship between dispersion and generation time. Previous studies find generation time to be unrelated to dispersion (Coale 1972; Tuljapurkar et al. 2009) but are they indeed independent? Based on our first finding, we hypothesized that dispersion *σ* is positively correlated to generation time *T_c_*. This hypothesis is indeed supported by our analyses, and we find that *σ* is proportional to *T_c_* on a loglog scale across taxa, and the explanatory power of this correlation is high. One basis for this result is previous work that suggests life-history traits such as generation time, age of maturity, lifespan-that scale as biological units of time (same dimension as time) have similar allometric exponents (close to 0.25) (Lindstedt and Calder III 1981). This result may also suggest that counter-examples of long-lived semalparous species (e.g., some salmonids) are rare.

We close by discussing the extensions of known allometric scaling relationships between generation time, intrinsic population growth rate, and average adult body mass, *M*. Previous studies have shown that generation time scales with body mass as *M*^0.25^ whereas density-independent intrinsic population growth rate scales with body mass as *M*^-0.25^(Charnov 1993; Brown et al. 2004). Our results imply that demographic dispersion scales with body mass in the same way as generation time. Further, our results suggest that the scaling of population growth rate with body mass holds in a wide range of environments, because the populations in our data are likely to be affected by density-dependence.

## Reproduction, Dispersion and Damping

Analyses of demographic damping have focused on humans or other species that can be described using age-structure (Keyfitz and Caswell 2005; Coale 1972; Trussell et al. 1977). In age-structured populations (in a constant environment), a life history is described by age-specific survival and reproduction. At age *x*, the average fertility is denoted by *m*(*x*) and the probability of surviving to age *x* by *l*(*x*). The expected lifetime reproduction of a newborn is the net reproductive rate *R_0_= ∑_x_ l*(*x*)*m*(*x*). For a cohort (individuals born at the same time), the generation time is the average age of reproduction

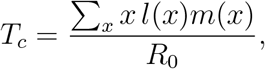

And the spread of reproduction around the mean age *T_c_* is captured by demographic dispersion *σ*, defined by

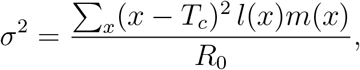

Similar expressions apply to stage-structured populations. In the paper by Steiner et al. (2014), the expressions for *T_c_* and *σ* (denoted as *V_a_* in their paper) were derived for stage-structured populations and have been used in this paper. Note that none of our data sets had both stages and ages, but for such cases there are appropriate formulas in Steiner et al. (2014).

A (st)age-structured population in discrete time is described by a population projection matrix, whose dominant eigenvalue is λ_0_= exp(*r*_0_) where *r*_0_ is the well-known intrinsic population growth rate. For the same matrix, the leading subdominant eigenvalue is in general complex λ_1_= exp(*r*_1_ + *is*_1_) (where *r*_1_, *s*_1_ are the real and imaginary parts of the sub-dominant root respectively and 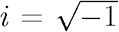). Here *r*_0_ should always be larger than *r*_1_. These eigenvalues are used to define the damping time *τ* and the damping rate *d*

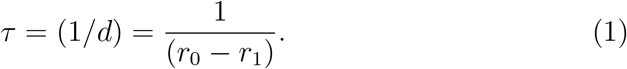

For each case we obtain a population projection matrix from the data (about which more below) and compute exactly (by standard numerical methods) the corresponding eigenvalues and the damping time.

After a disturbance, the population structure changes as the product *e^−dt^* cos (2*πt/T*), with cycles of period *T ⋍ T_c_* whose amplitude decreases at damping rate *d* > 0. Here *d* = (*r*_0_—*r*_1_), and *t* is time since the disturbance. Damping with *d* > 0 is assured because in general *r*_0_> *r*_1_ for (st)age structured models (Caswell 2001). Thus the damping time in equation (1) is the time scale of convergence to the stable (st)age distribution.

Our work was motivated by Wachter (1991) extending earlier work on age-structured populations. He used the Lotka renewal equation to approximate *r*_0_, *r*_1_, *s*_1_. For small growth rates,

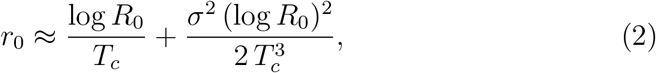

whereas

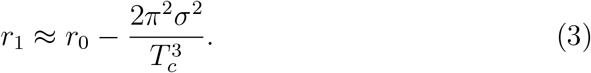

A similar expression for *r*_0_ holds in stage-structured populations (Steiner et al. 2014). In such cases, we conjecture that *r*_1_ is also given by the approximation equation (3).

Using these approximations in equation (1), the damping time is

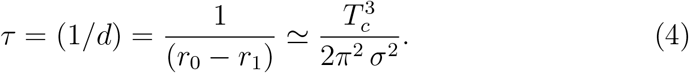

Hence we expect that damping time *τ* should increase with generation time *T_c_*, and decrease with increasing age dispersion *σ* of reproduction.

## Results

### Damping time and generation time

From equation (4) we expect that damping time (calculated from the population projection matrix) should increase with both generation time and demographic dispersion. However, we find that damping time is positively correlated with generation time across 664 species of animals and plants (on a logarithmic scale, see Fig 1), regardless of demographic dispersion. The relationship is significant but noisy, more so for stage-based population models than age-based ones. The variability around the main correlation that is evident in Fig 1 may have several sources:

a. the expression (4) is an approximation so higher moments of the distribution of reproduction may be significant for some species, especially those with stage-based dynamics;
b. the data reflects sampling variability, especially for populations with small population size in study, and so some variation is to be expected;
c. in some populations that have long-lived stage(s), such as trees, the numbers of deaths to large individuals observed during the study period may be small so the corresponding estimated survival rates may be artificially high.

**Figure 1:**
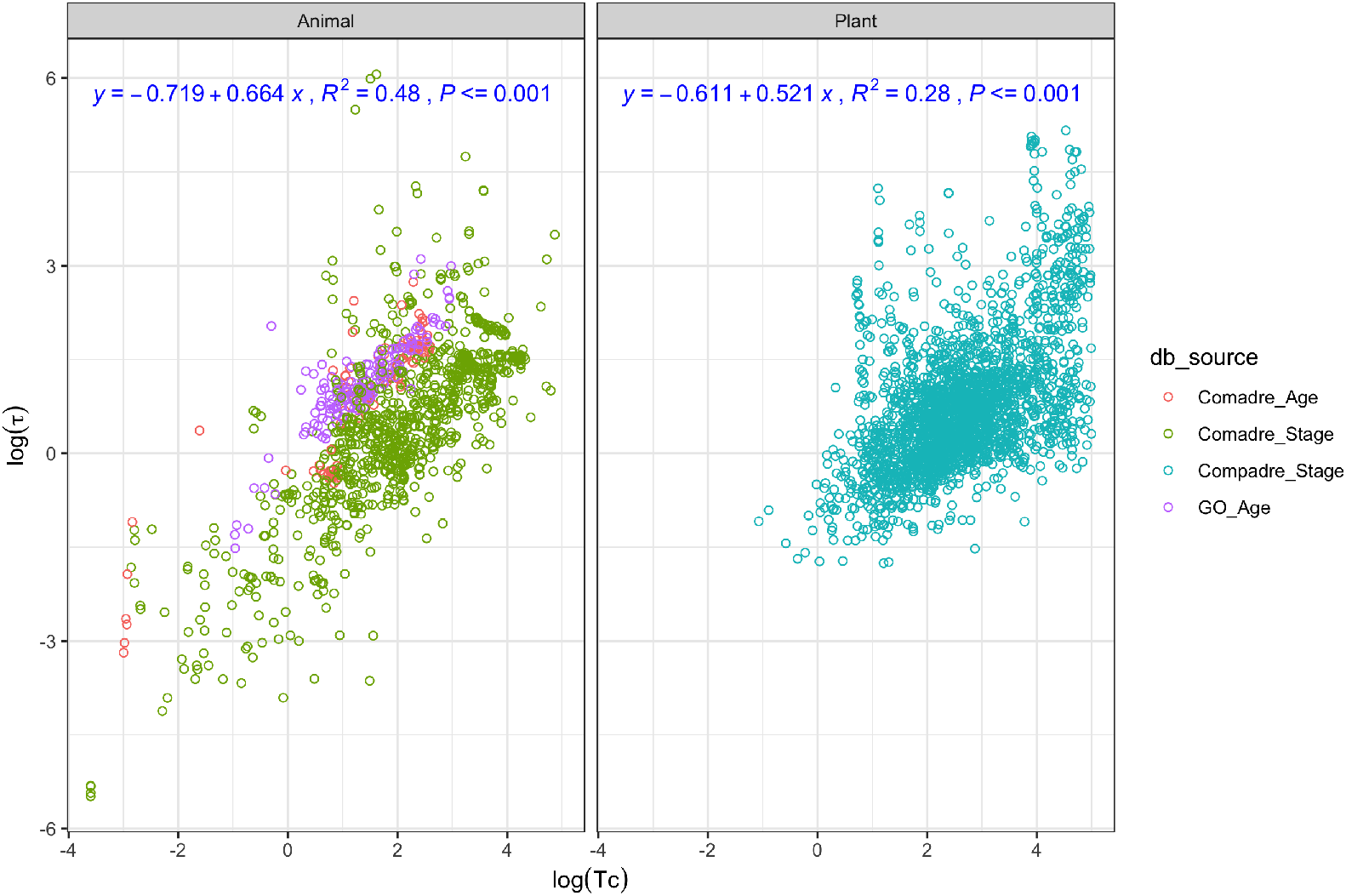
Damping time (*τ*) versus generation time (*T_c_*) on a log-log scale for animals (*left* panel) and plants (*right* panel). The colors correspond to data as in the previous figure. The top left of each panel presents the fitted model, its coefficient of determination (*R*^2^) and P-value based on linear regression. The standard error of the regression slope is 0.019 (left panel) and 0.017 (right panel). The damping time (*τ*) used is calculated exactly from each population projection matrix. The regression slopes for each dataset are: Comadre_Age (0.81), Comadre_Stage (0.72), Compadre_Stage (0.52) and GO_Age (0.76).

Even so, Fig 1 clearly supports the conclusion that species with short generation time can recover rapidly from environmental disturbances and are less vulnerable to such perturbations. On the other hand, species with long generation time are more vulnerable as they take longer to converge to their stable (st)age demographic distribution. Across species we find a scaling relationship

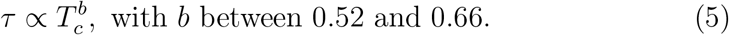

Drilling down, we asked if a similar relationship between damping time and generation time hold for the species within biological classes? We found a positive relationship to hold within most classes, though the variability is large (see Appendix figures B.1 and B.2). The relationship is stronger in classes that contain a large number of species in our data, such as *Actinoptery-gii, Aves, Mammalia*, and *Reptilia* in animals and *Magnoliopsida* and *Liliop-sida* in plants.

### Generation time and demographic dispersion

Based on the first result, we hypothesize *T_c_* is proportional to *σ* given the approximation (See equation (4)). Our hypothesis is supported as the result indicates generation time and dispersion to be positively related, evidenced by the remarkable linear correlation (on a logarithmic scale, see Fig 2) between the two. The relationship between *T_c_* and *σ* is statistically significant and the degree of explanation is high with an *R*^2^ statistic of 0.95. The regression slope of log(*σ*) versus log(*T_c_*) is close to 1 for a wide range of taxonomic groups in both plants and animals. Hence we find a scaling relationship:

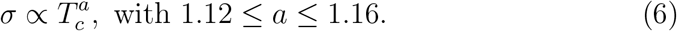

**Figure 2:**
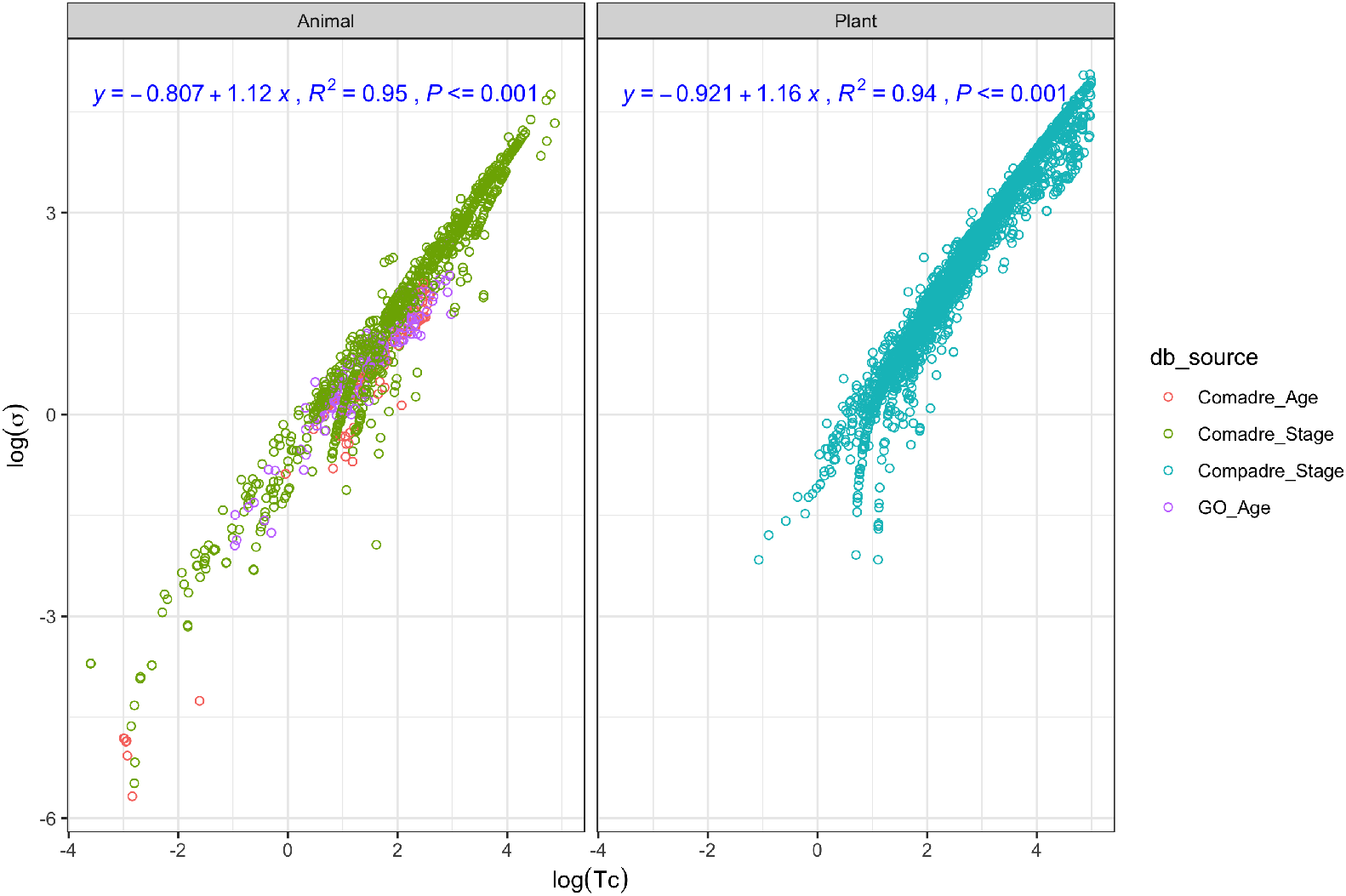
Demographic dispersion (*σ*) versus generation time (*T_c_*) on a log-log scale for animals (*left* panel) and plants (*right* panel). Age-structured animal data from Comadre (*left*, red), stage-structured animal data from Comadre (*left*, grass green), age-structured mammal data from GO (*left*, purple) and stage-structured plant data from Comapdre (*right*, lake blue). The top left of each panel presents the fitted model, its coefficient of determination (*R*^2^) and P-value based on linear regression. The standard error of the regression slope is 0.007 (left panel) and 0.006 (right panel). The regression slopes for each dataset are: Comadre_Age (1.20), Comadre_Stage (1.11), Compadre_Stage (1.16) and GO_Age (0.93).

To analyze this scaling in more detail, we grouped species by biological classes, and within each class found a strong positive (logarithmic) relationship between demographic dispersion and generation time that contain a large number of species in our data. These classes include *Actinopterygii, Aves, Mammalia*, and *Reptilia* in animals and *Magnoliopsida, Phaeophyceae* and *Liliopsida* in plants. The regression slope between log *T_c_* and log *σ* for each Class varied but were approximately close to 1 (see Appendix figures B.3 and B.4).

At the opposite extreme of biological detail, we can argue that in an age-structured population an increase in demographic dispersion *σ* likely implies a larger reproductive span as measured by the difference between age at last reproduction *ω* and age at first reproduction *α*. Based on our result for *σ*, a simple hypothesis is that reproductive span (*ω – α*) also increases with generation time, and indeed we found such a positive relationship, although noisier (see Appendix figure B.5).

### Allometric scaling of life history measures with body mass

We extend our results through allometric scaling relationships as life-history variables such as-age of first reproduction, longevity, adult mortality, generation time and maximum intrinsic population growth rate, have been shown to scale with average adult body mass (Blueweiss et al. 1978; Read and Harvey 1989; Promislow and Harvey 1990; Gillooly et al. 2002; Brown et al. 2004; Hamilton et al. 2011; Hatton et al. 2019). An immediate consequence of our results is that any allometric scaling for *T_c_* must imply a corresponding scaling for *σ* and *τ*.

Empirically, the generation time, *T_c_*, scales with average adult body mass *M* (Millar and Zammuto 1983; Gillooly et al. 2002; Brown et al. 2004; Hamilton et al. 2011; Brown et al. 2018), as

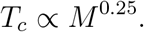

Using the above scaling in equation (6) with *a* ⋍ 1 we find:

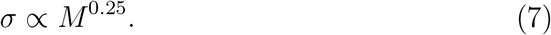

From equation (4), the damping time scales approximately as

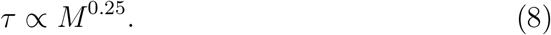

Next use equation (6) but with *a* ⋍ 1 in the growth rate equation (2) to find

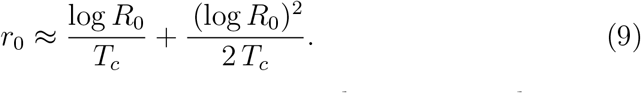

The net reproductive rate, *R*_0_, is not expected to vary with average adult body mass *M*, based on theoretical arguments of Brown et al. (2018), Charnov (1993), and Pianka (1988), and empirical studies in the absence of density-dependent feedbacks (Charnov et al. 2007; Ginzburg et al. 2010). Even supposing that *R*_0_ has an allometric dependence on *M*, the quantity log *R*_0_ in equation (9) would vary with log *M* and slowly with *M*. Given such a weak dependence, equation (9) implies that the relationship between *r*_0_ and *M* is mainly due to (1/*T_c_*), and hence

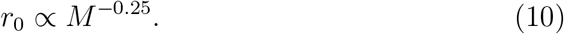

This scaling was previously known for growth rate without density-dependence (Hennemann 1983; McMahon and Bonner 1983; Charnov 1993) (i.e., for the maximum intrinsic population growth rate). In general, empirical data to estimate *r*_0_ is collected across a wide range of density conditions. Given the diversity of environmental contexts, we did not expect observed *r*_0_ to display the above allometric relationship. However, we think large number of matrices in our dataset may offset the effect of density conditions.

## Discussion

Even though not all populations converge to a stable (st)age distribution in the real world, the patterns of convergence to and deviation from the stable (st)age distribution can still provide useful biological information (Carey 1983; Keyfitz and Caswell 2005). Our first finding reveals a robust positive relationship between damping time and generation time. We find damping time (inverse of damping rate) is positively correlated with generation time on the log-log scale, regardless of demographic dispersion. This implies that species with slow life-histories (characterized with long generation time, late maturity, low fecundity) are more vulnerable to negative environmental disturbances than species with fast life-histories. This finding is supported by Capdevila et al. (2020) who have argued that the long convergence time (slow recovery rate) of the Asian elephant (*Elephas maximus*) makes them more vulnerable to continuous habitat loss than the red squirrel (*Tamiasci-urus hudsonicus*). Lebreton et al. (2012) also find that conservation status deteriorates with increasing generation time. Di Camillo and Cerrano (2015) documented mass mortality events in the North Adriatic sea and observed a shift of the benthic assemblage from slow-growing and long-lived species (of the sponge *C. reniformis*) to a community dominated by fast-growing and short-lived cnidarians. Previous studies on Amazonian mammals (Bodmer et al. 1997), and birds (Owens and Bennett 2000) indicate that species with long generation times are more prone to the threat of extinction. Therefore, generation time, which is not only a major axis of variation in life-history tactics in mammals (Gaillard et al. 1989), also sets the time scale for response and recovery of species. Our study of recovery times from disturbances can inform conservation planning (Salguero-Gómez et al. 2016).

Our first result led us to the finding that demographic dispersion and generation time are positively correlated on the log-log scale. This result implies that long-lived (i.e., slow) species spread reproduction over a wider age range than the converse. One possible explanation is that the large demographic dispersion enables slow-lived species (characterized by low fecundity) to increase offspring survival by investing more energy in each reproductive event. Also, species with slow life histories may take a long time to reach the reproductive window, and so experience large variability during their development (e.g. differences in resource utilization, growth in body size). Such variability has a modest effect on fitness in long-lived populations (Tuljapurkar 1982), and can ride out fluctuations by averaging over several ages (Sæther et al. 2013). In addition, previous theoretical arguments (Tuljapurkar 1982) have shown that fitness as defined by stochastic growth rate depends on intrinsic growth rate *r*_0_ which is a function of *σ* and *T_c_* (Wachter 1991). Therefore, our result on the correlation between *σ* and *T_c_* has implications for stochastic growth rate and population dynamics in fluctuating environments.

Our results also suggest several allometric scaling relationships between average body mass and the life history parameters of demographic dispersion, damping time, and intrinsic population growth rate. The allometric scaling of dispersion we found contributes to previous work that suggests life-history traits that scale as biological units of time have similar allometric exponents (close to 0.25) (Lindstedt and Calder III 1981). Our result on growth rate also extends the previously established metabolic scaling from a density-independent scenario (Charnov 1993) to a wide range of environments. The metabolic theory of ecology (West et al. 1997, 1999; Brown et al. 2004) may explain such empirical observations in terms of basic biological and physiological processes.

There are limitations in our study that call for future research. Considering our dataset spans a wide range of species, we did not analyse phylogenetic relationship between species which may provide scope for further work. Our analyses use the theory of density-independent time-invariant population projection models, whereas research that takes density-dependence and stochas-ticity into account is needed to provide a more comprehensive picture of life-history and transient dynamics. We only consider recovery from a single disturbance event and have not studied the effects of disturbances of varying magnitude or duration that may impact populations through different mechanisms (Owens and Bennett 2000). Our arguments about life-history strategies and the slow-fast continuum focus on animals (largely mammals). For plants, we need a better understanding of their life-histories to explain our findings. A deeper understanding of the deviations we observe in our results may require a more searching analysis of individual life-histories.

## Data

Our aim is to examine damping time for a large number of species covering a wide range of generation time. To do so we used three databases which provide population projection matrices: COMPADRE (v.6.20.5.0) for plants (Salguero-Gómez et al. 2016); COMADRE (v.4.20.5.0) for animals (Salguero-Gómez et al. 2016); and compiled lifetables for mammals by Jean-Michel Gaillard (Gaillard et al. 2005) and Madan Oli (Oli 2004) (hereafter GO).

After data checking and cleaning (details in the Appendix), we have 3622 matrices (664 different species) in total: in COMPADRE there are 2353 stage-structured matrices (331 species); in COMADRE, there are 1029 stage-structured matrices (217 species) and 96 age-structured matrices (32 species). In the GO dataset, there are 144 age-structured matrices (112 species) after removing the matrices that were also present in COMADRE dataset.

## Appendix A Supplementary Information on Data

For COMPADRE (v.6.20.5.0) and COMADRE (v.4.20.5.0) database, there were initially 8925 matrices (759 species) and 2275 matrices (415 species), respectively. Then we conducted a series of data cleaning to prepare the dataset for the analysis.

### A.1 Data classification for age- and stage-structured matrices

The database consists of three criteria to indicate whether the population projection matrix contains (st)ages based on size (MatrixCriteriaSize), development (MatrixCriteriaOntogeny), age (MatrixCriteriaAge). For COMADRE and COMPADRE, we first take out “NA”s in these three criteria, then use them to get a rough classification of age and stage-structured data: if Ma-trixCriteriaSize ==“No” & MatrixCriteriaAge ==“Yes” & MatrixCrite-riaOntogeny ==“No”, it’s considered as age-structured data, otherwise it’s stage-structured data.

For COMPADRE, we only consider stage-structured data for the analysis. For age-structured data in COMADRE, we further check the the intersection of last row and last column in the survival matrix. If the value is zero, then we classify it as a age-structured data; if it is non-zero, then classify it as stage-structured data considering the population is alive in the last stage observed.

### A.2 Filters used before calculation

Using the flags in the dataset, we exclude data with missing values in vital rates (i.e, to ensure no NA’s in the matrices); ensure the ergodicity of population projection matrix; ensure the survival for a given age/stage is always less than or equal to 1; ensure the fecundity was measured in the study; eliminate those matrices with non-zero cloning data; keep matrices where the projection interval is non-zero; remove data from one unclear source (Master thesis) with no title and author name.

We also remove semelparous species *Oncorhynchus tshawytscha* (Chinook salmon); ensure the fertility matrix has elements only on the first row and the rest of the rows are all 0; remove data with males and females vital rates separately in the same population projection matrix; remove Bacteria (*Spirochaetes*) and Virus (*lentivirus*); remove data with the intersection of last row and last column of survival matrix equals to 1.

For age-structured data in COMADRE, we further ensure survival matrix should only have non-zero value in sub-diagonal; ensure that the fertility matrix has more than 1 non-zero value in the first row.

### A.3 Filters used during and after calculation

For stage-structured data in both COMADRE and COMPADRE, we remove matrices where (I-U) inverse does not exist (where I is the Identity matrix and U is the survival matrix) to enable the calculation. It should be noted that we standardized the databases so that the projection interval is the same (1 year) for all species. Besides, considering the biological realisticity, we removed unlikely values by ensuring log(*T_c_*) < 5, log(*σ*) >-15, and log(*τ*) < 15.

### B Figure

**Figure B.1:**
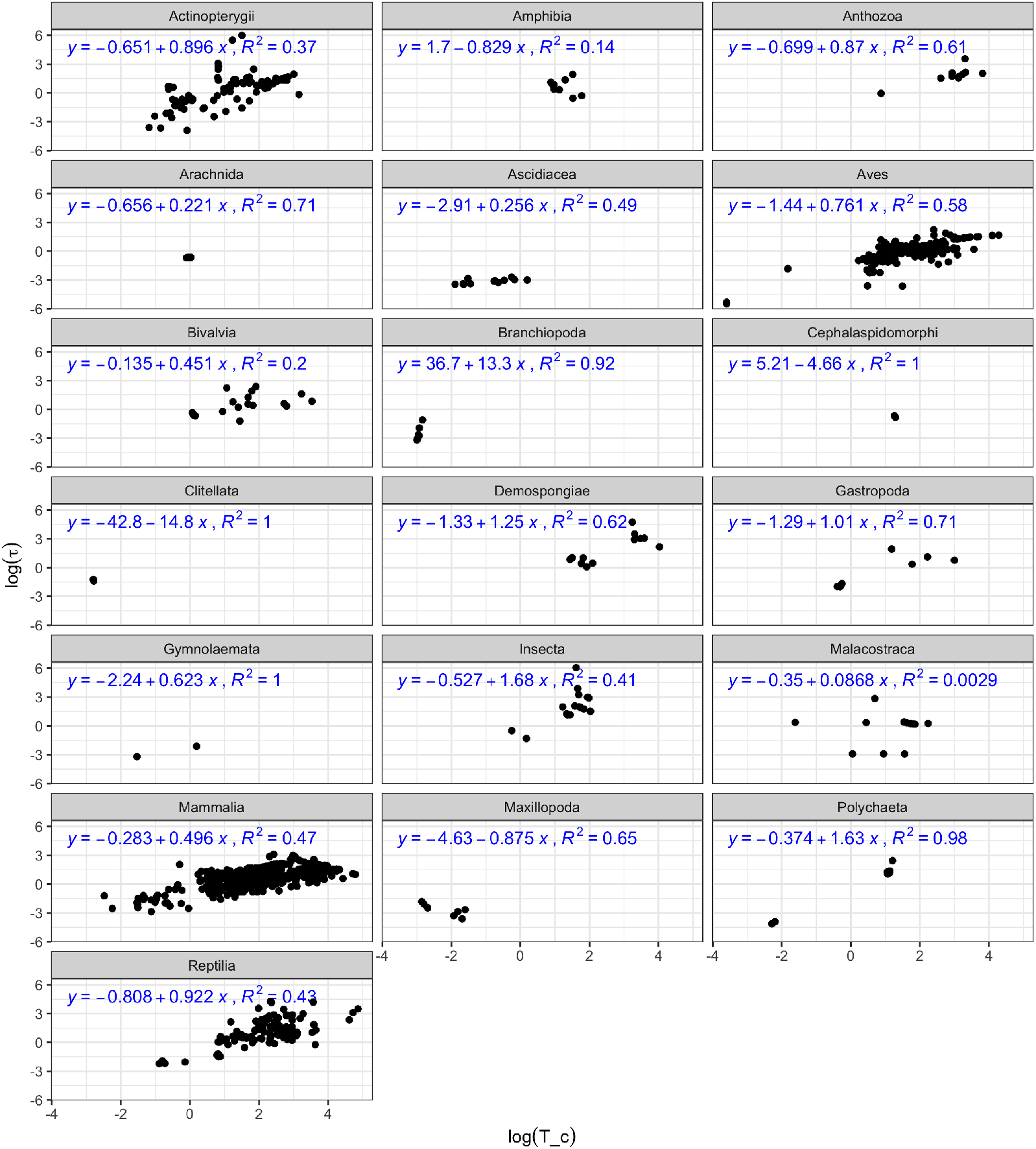
Class-wise plots for damping time (*τ*) versus generation time (*T_c_*) on a log scale for animals. Each panel corresponds to a Class. On the top left of each panel, we also present the fitted model and its coefficient of determination (*R*^2^) based on linear regression. It should be noted that the damping time (*τ*) presented here is the exact value calculated from population projection matrix instead of the approximation in equation (4).

**Figure B.2:**
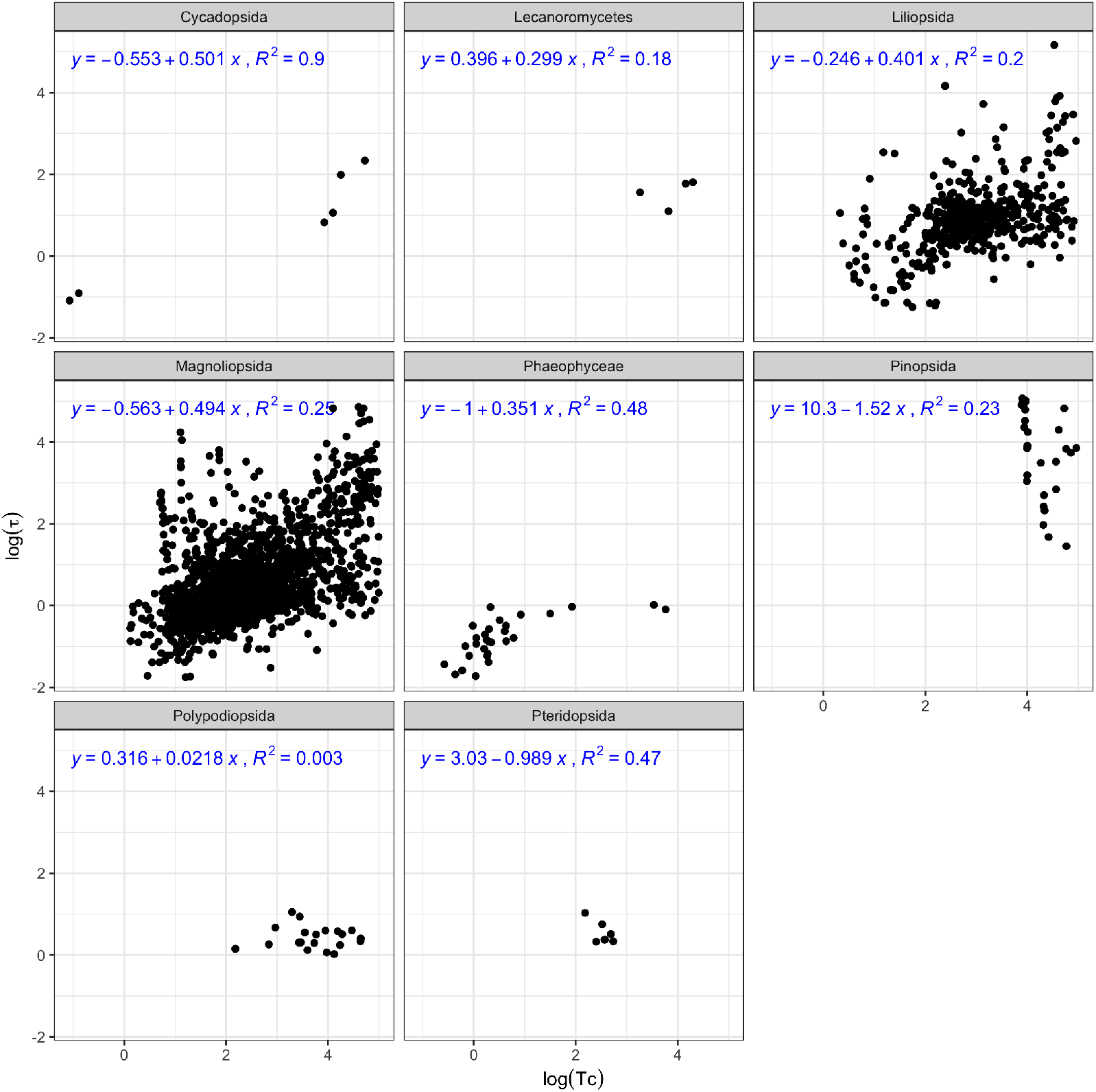
Class-wise plots for damping time (*τ*) versus generation time (*T_c_*) on a log scale for plants. Each panel corresponds to a Class. On the top left of each panel, we also present the fitted model and its coefficient of determination (*R*^2^) based on linear regression. It should be noted that the damping time (*τ*) presented here is the exact value calculated from population projection matrix instead of the approximation in equation (4).

**Figure B.3:**
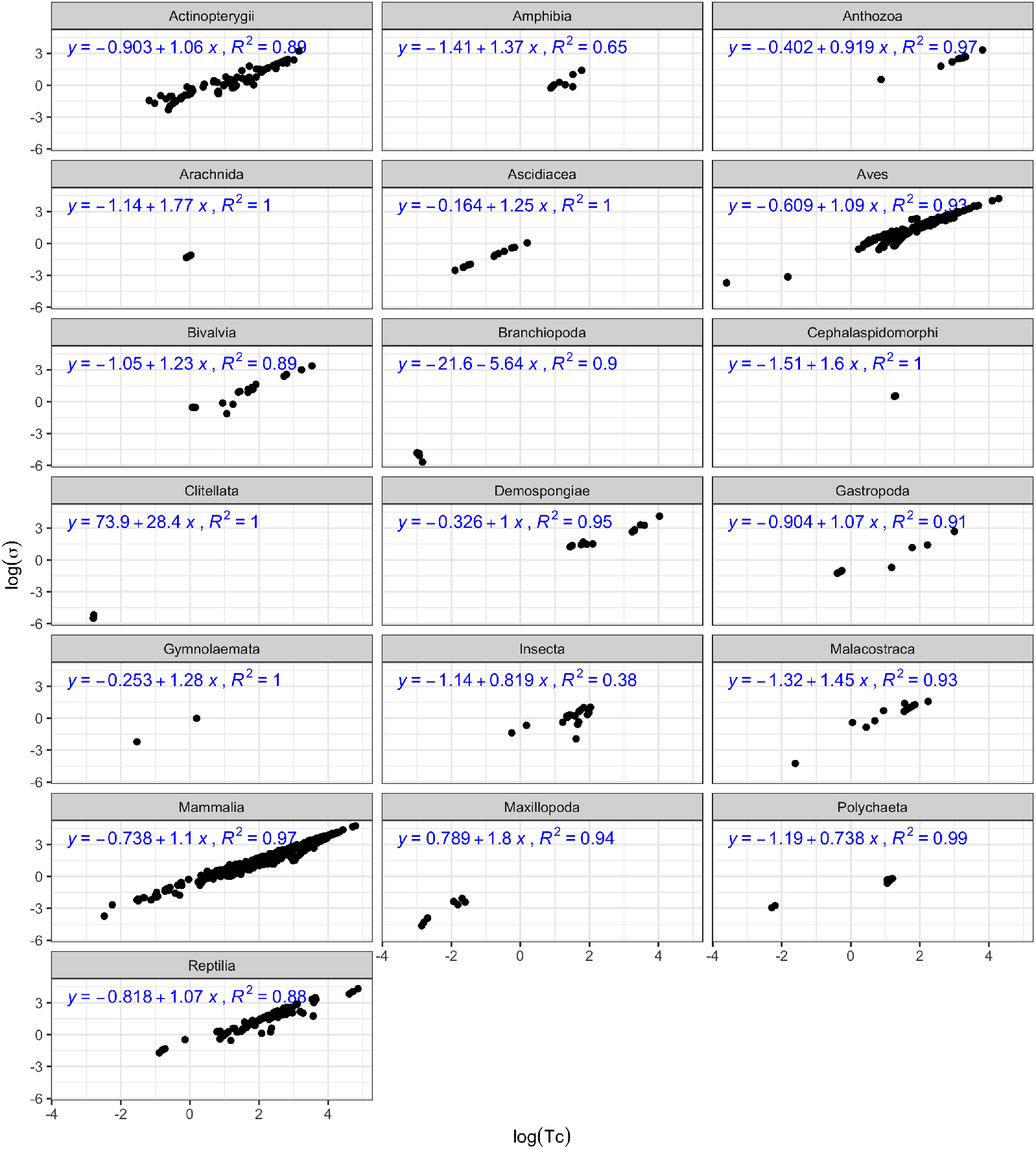
Class-wise plots for demographic dispersion (*σ*) versus generation time (*T_c_*) on a log scale for animals. Each panel corresponds to a Class. On the top left of each panel, we also present the fitted model and its coefficient of determination (*R*^2^) based on linear regression.

**Figure B.4:**
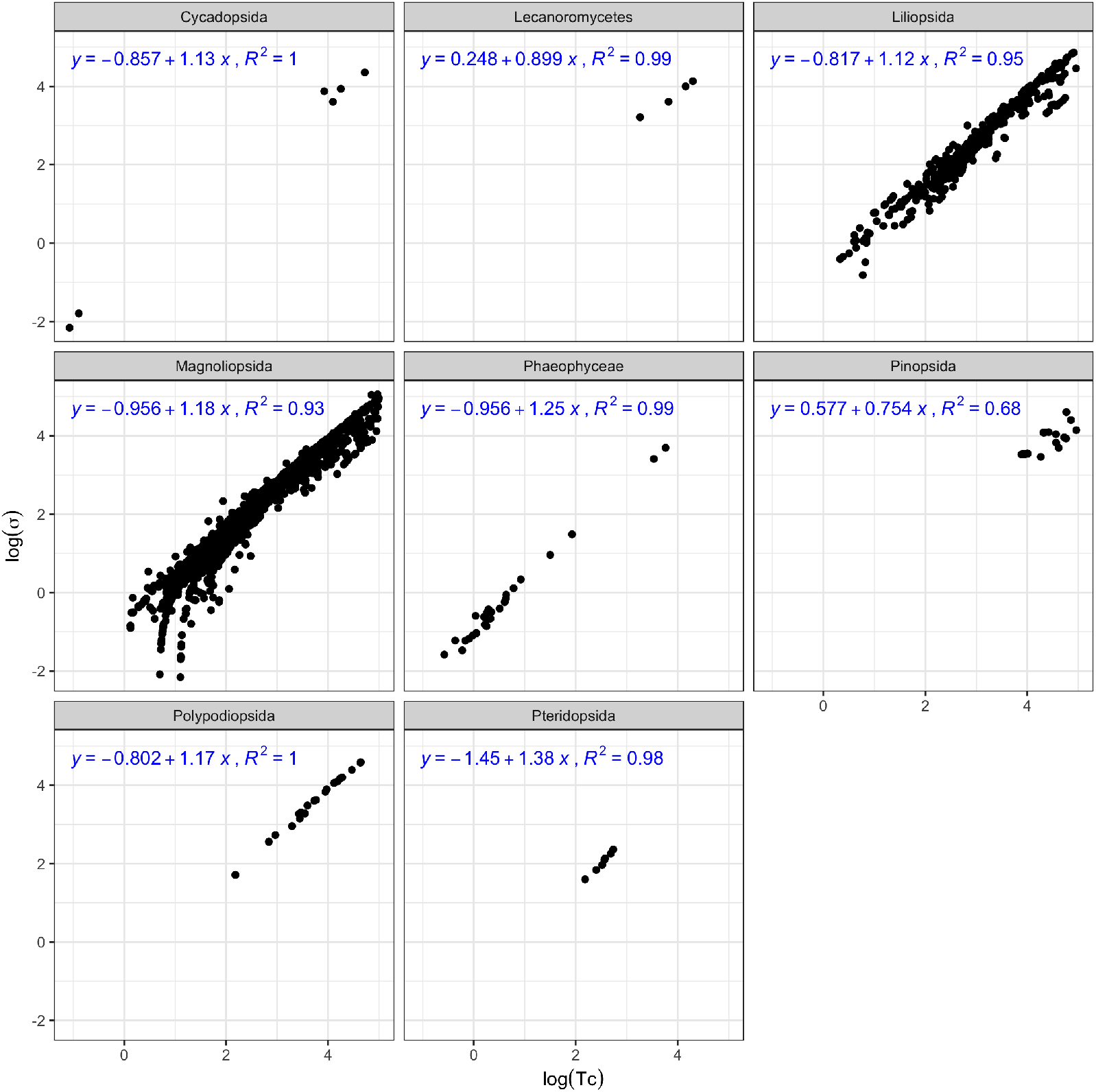
Class-wise plots for demographic dispersion *σ* versus generation time *T_c_* on a log scale for plants. Each panel corresponds to a Class. On the top left of each panel, we also present the fitted model and its coefficient of determination (*R*^2^) based on linear regression.

**Figure B.5:**
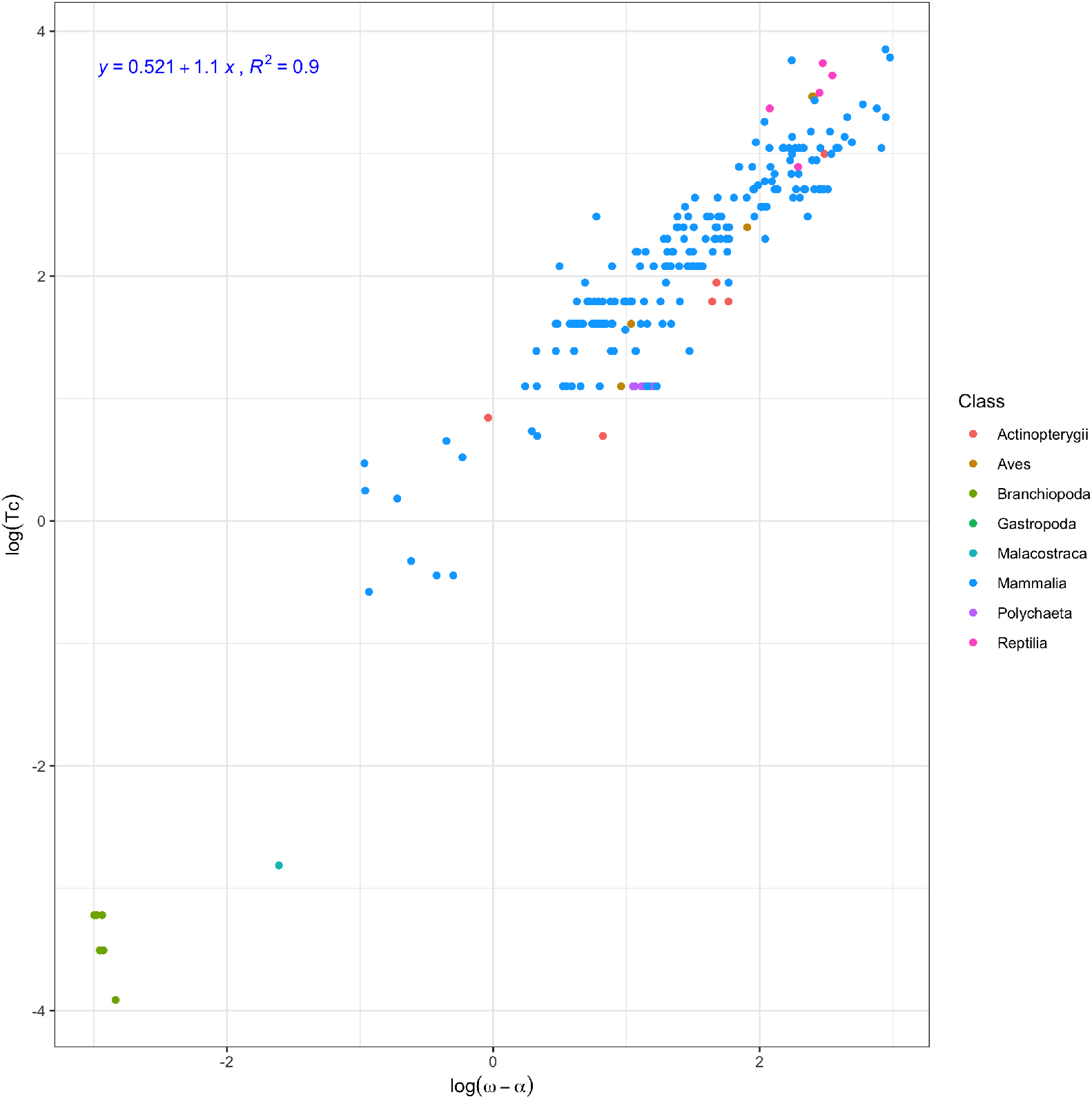
Reproductive span (*ω* – *α*) versus generation time (*T_c_*) on a log scale for age-structured animals. Different colors indicate different biological Classes. On the top left, we also present the fitted model and its coefficient of determination (*R*^2^) based on linear regression.

